# Repeated presentation of visual threats drives innate fear habituation and is modulated by environmental and physiological factors

**DOI:** 10.1101/2024.10.15.618513

**Authors:** Jordan N. Carroll, Brent Myers, Christopher E. Vaaga

## Abstract

To survive predation, animals must be able to detect and appropriately respond to predator threats in their environment. Such defensive behaviors are thought to utilize hard-wired neural circuits for threat detection, sensorimotor integration, and execution of ethologically relevant behaviors. Despite being hard-wired, defensive behaviors (i.e. fear responses) are not fixed, but rather show remarkable flexibility, suggesting that extrinsic factors such as threat history, environmental contexts, and physiological state may alter innate defensive behavioral responses. The goal of the present study was to examine how extrinsic and intrinsic factors influence innate defensive behaviors in response to visual threats. In the absence of a protective shelter, our results indicate that mice showed robust freezing behavior following both looming (proximal) and sweeping (distal) threats, with increased behavioral vigor in response to looming stimuli, which represent a higher threat imminence. Repeated presentation of looming or sweeping stimuli at short inter-trial intervals resulted in robust habituation of freezing, which was accelerated at longer inter-trial intervals, regardless of contextual cues. Finally, physiological factors such as acute stress further disrupted innate freezing habituation, resulting in a delayed habituation phenotype, consistent with a heightened fear state. Together, our results indicate that extrinsic factors such as threat history, environmental familiarity, and physiological stressors have robust and diverse effects on defensive behaviors, highlighting the behavioral flexibility in how mice respond to predator threats.

## Introduction

The ability of animals to accurately and appropriately respond to predator threats in the environment is critical for survival. As such, antipredator defensive behaviors are observed across evolutionary history (D. C. Blanchard & Blanchard, 2008; Kavaliers & Choleris, 2001; Kikuchi et al., 2023; LeDoux, 2012). Such antipredator defensive behaviors form the basis of unconditioned or innate fear responses, which are distinct from conditioned fear, in that they do not require previous associative learning; suggesting the presence of hard-wired, dedicated neural circuitry for the detection, integration, and execution of appropriate behavioral responses (Carrive, 1993; De Franceschi et al., 2016; Evans et al., 2018; Keay & Bandler, 2001; LeDoux, 2012; B. A. Silva et al., 2016; Yilmaz & Meister, 2013; Zhang et al., 1990). For example, mice show a dynamic repertoire of defensive behaviors which are differentially engaged depending on the nature of the threat (De Franceschi et al., 2016; Fanselow, 1991, 1994; Tafreshiha et al., 2021). These observations have informed the development of the threat imminence model, in which specific sensory stimuli are ethologically matched to appropriate defensive behaviors (Bolles, 1970; Fanselow, 2018; Fanselow & Lester, 1988; Perusini & Fanselow, 2015). For instance, sweeping visual stimuli that mimic a distal aerial predator engage freezing behaviors to avoid detection. Conversely, looming visual stimuli, which mimic a proximal aerial threat, engage more active defensive strategies such as flight to a shelter (De Franceschi et al., 2016; Liu et al., 2022; Solomon et al., 2023; Yilmaz & Meister, 2013)

Consistent with the threat imminence theory, freezing and flight behaviors are thought to be differentially engaged by distinct rostro-caudal columns in the midbrain periaqueductal gray (Bandler et al., 1985, 2000; Bandler & Shipley, 1994; Carrive, 1993; Keay & Bandler, 2001; Tovote et al., 2016; Zhang et al., 1990). More specifically, activation of the ventrolateral column of the periaqueductal gray (vlPAG) results in robust freezing behaviors, whereas activation of the dorsolateral periaqueductal gray (dlPAG) results in active avoidance strategies, such as flight (Bandler & Shipley, 1994; Carrive, 1993; La-Vu et al., 2022; Tovote et al., 2016; Vaaga et al., 2020; Zhang et al., 1990). Despite this theoretical and neural framework, innate fear behaviors are not fixed responses, and therefore may be modulated by environmental and physiological variables. For example, looming threats can elicit *freezing* in experimental conditions without a protective shelter (De Franceschi et al., 2016; Yilmaz & Meister, 2013).

This observation raises the possibility that other environmental factors, such as threat history, environmental familiarity, or physiological factors, such as exposure to acute stress, may similarly alter innate fear responses (Hassien et al., 2020; Lenzi et al., 2022; Perusini & Fanselow, 2015; Rau et al., 2005; Tafreshiha et al., 2021). However, one limitation of the threat imminence model is that behavioral variables such as response vigor are often inferred by the defensive strategy employed, limiting direct comparisons. As such, understanding how environmental and physiological variables impact innate fear behavior has been difficult to assess. Of particular interest is how such variables contribute to behavioral flexibility, as inflexible fear responses are observed in disorders such as post-traumatic stress disorder (PTSD; Friedman et al., 2011; Iqbal et al., 2023; Koenen et al., 2017)

To begin to understand how intrinsic and extrinsic factors influence innate fear and behavioral flexibility, we exposed mice to looming and sweeping threats in an arena without a shelter, in an attempt to limit the available defensive behavioral repertoire. We demonstrate that under such conditions, both sweeping and looming threats engage immobility behavior, although threat imminence is still encoded by response vigor (i.e. freezing duration). Repeated threat presentation resulted in a progressive reduction in immobility (see also (Lenzi et al., 2022), independent of the nature of the visual stimulus. Furthermore, the rate, but not degree, of habituation significantly varied with changes in environmental condition and/or physiological stressors, suggesting that threat habituation is a key variable in the innate fear response.

## Methods

### Ethical Note

All experimental procedures were conducted in accordance with institutional guidelines regarding the ethical use of animals. All experimental methods were approved by Northwestern University (protocol IS00014844, IMR) and Colorado State University Institutional Animal Care and Use Committees (protocol 3836, CEV). The study utilized (6-12 week) adult wild-type mice, purchased from a commercial supplier (Jackson Laboratories). Experiments involved non-invasive behavioral observations of animals exposed to visual stimuli mimicking predators. All reasonable efforts were made to increase scientific transparency and openness. All original data, python code, and digital research materials are available upon reasonable request. The study design and analysis were not pre-registered.

### Animals

Adult male and female C57Bl6/J wild type mice were used for the study, in sex balanced cohorts. Cohorts of 10 wildtype mice (5 male and 5 female) were purchased from Jackson Laboratories (Strain: 000664) at 4-6 weeks of age and allowed to recover from transport stress in the animal facility for at least 2 weeks prior to behavioral testing. Mice were socially housed (2-5 mice per cage) on a 12:12 hour light:dark cycle with *ad libitum* access to food and water.

At least 1 week prior to behavioral testing, mice were transported to a satellite housing facility located in the same building as the behavioral testing suite to reduce daily transport stress. Much of the data was collected during the animal light cycle, although in a subset of cohorts, testing was performed during the animal dark cycle. No differences in behavior were observed across the light cycle, so data were pooled. To reduce potential circadian effects on arousal and behavioral responses, animals tested during the light cycle were allowed to acclimate to the dark behavioral testing suite for at least 30 minutes prior to testing. Unless otherwise noted, two days prior to behavioral testing, all animals underwent at least 2 days of handling and behavioral familiarization in the experimental chamber for at least 10 minutes each day. Following familiarization trials, mice were placed in a temporary holding cage before all mice were returned to their home cage.

*Innate Fear Paradigm:* The experimental setup consisted of a 25 × 25 × 25 cm acrylic behavioral chamber with 3 light grey walls and 1 transparent wall to facilitate video recordings of animal behavior. Visual stimuli were presented using an LCD monitor placed 40 cm above the arena floor. The sweeping visual stimulus (Figure 1B,left) consisted of a high contrast disk (5° visual angle) which traversed the screen and returned to its original position in a total of 8 seconds (De Franceschi et al., 2016). The looming visual stimulus (Figure 1B,right) consisted of a high contrast rapidly expanding disk which expanded to 20° visual angle in 333 ms and repeated 5 times in a total of 6 seconds (Yilmaz & Meister, 2013).

**Figure 1:**
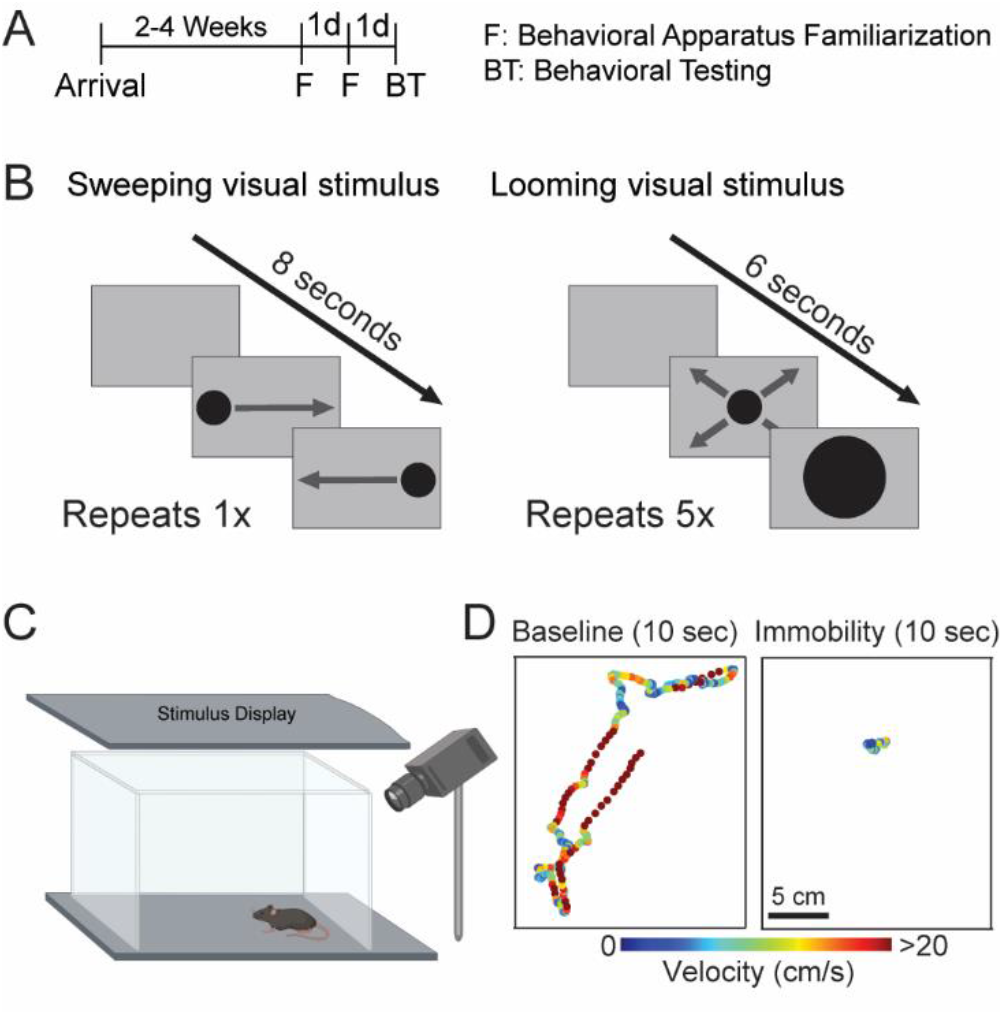
Experimental Methodology. (A) Timeline of typical experiment. Mice were familiarized with the behavioral arena for at least two days prior to behavioral testing. (B) Schematics illustrating the sweeping (left) and looming (right) visual stimuli used to elicit innate fear responses. (C) Diagram of the behavioral chamber, indicating relative position of the stimulus display and video camera. (D) Exemplar output from DeepLabCut illustrating average animal position within the behavioral chamber during the baseline period and the 10 seconds immediately after presentation of a looming stimulus. The data point from each frame is pseudo-colored by velocity.

Both the stimulus presentation and video acquisition were controlled using custom-written python modules and a raspberry pi system. Briefly, videos were recorded using an infrared raspberry pi camera module at 15 or 30 fps. Because the recordings were done in the dark, the only ambient light was from the overhead monitor. Additional infrared lights were used to evenly illuminate the behavioral arena. Visual stimuli were manually triggered using a Master-8 programmable pulse generator. Stimuli were triggered after 2-3 minutes of baseline activity to allow the mice to refamiliarize themselves with the behavioral chamber. Furthermore, attempts were made to trigger the stimuli during periods of movement, to adequately capture freezing behaviors. For most experiments, mice were exposed to three identical stimuli in a single behavioral session, separated by an inter-trial interval of ∼5 minutes. In some experiments, the inter-trial interval was increased to 24 hours, to test longer term behavioral habituation.

### Acute Stress Paradigm

To test for effects of acute stress on innate fear, mice were exposed to a modified stress-enhanced fear learning paradigm (Hassien et al., 2020; Perusini et al., 2016; Rau et al., 2005; Rau & Fanselow, 2009). After 2 days of familiarization in the open field arena, mice were placed in a novel context fear conditioning chamber to undergo an acute stress paradigm. After approximately 1 minute, mice were given 4 unconditioned, unpredictable foot shocks (2 sec duration, 1 mA) at an interval of 60-80 seconds (Hassien et al., 2020). Mice were then allowed to recover for either 1 hour or 24 hours before being placed in the open field arena for innate fear testing.

### Data Analysis

Videos were initially analyzed using DeepLabCut marker-less pose estimation to track animal position. Subsequent data analysis was performed using custom python code.

First, for each frame, the x-and y-position of the mouse’s center of mass was identified (Figure 1D). Animal speed was calculated frame-by-frame by dividing the change in animal position by the interval frame rate. Velocity data was smoothed using a rolling average across 10 frames, and an immobility (freezing) epoch was defined as any 500 ms period in which the animal velocity was less than 2 cm/sec. To facilitate data presentation, velocity traces were then filtered to only show periods of immobility. Percent immobility was calculated within the 20 second period after the onset of the visual stimulus.

### Statistical Testing

Data are reported as mean±S.E.M unless otherwise noted. Data analysis and statistical testing was performed in GraphPad Prism software. Statistical comparisons between two groups were calculated using either a two-sample paired or unpaired t-test, as indicated in the text. For experiments in which we compared immobility across trials, data was analyzed using a one-way repeated measures ANOVAs with a Tukey post-hoc comparison. Comparisons of normalized innate fear habituation were compared using a ordinary one-way repeated measures ANOVA. Datasets were compared using a two-way repeated measure ANOVA.

As stated above, all experiments were performed with sex-balanced cohorts, which, unless otherwise indicated, were pooled together for analysis. To further reduce bias, animals were randomly assigned to experimental cohorts. Data was analyzed using a pipeline to reduce experimenter bias. The n values reported reflect the number of animals in each experiment. Sample sizes were determined using a power analysis with preliminary data, which indicated that a sample size of 10 mice per group was sufficient to detect a biologically relevant effect size of ∼20% with a statistical power (β) of 0.8 and at a type I error rate (α) of 0.05. Exclusion criteria included low baseline movement, which would occlude the ability to detect freezing behavior; however, no animals were excluded from the dataset using this criterion. One animal was excluded, as described in the text, from further analysis because its response was greater than three times the standard deviation of the population response.

## Results

### Freezing responses to sweeping and looming visual stimuli

Under experimental conditions where animals have access to a shelter, looming (proximal) and sweeping (distal) threats engage distinct active and passive coping behavioral strategies, respectively (De Franceschi et al., 2016; Evans et al., 2018; Tafreshiha et al., 2021; Yilmaz & Meister, 2013). Such distinct behavioral responses to proximal vs. distal threats have precluded testing whether and how mice differentially encode threat imminence via changes in response vigor. We therefore sought to directly compare behavioral responses to looming and sweeping visual threats under conditions in which the available defensive strategies are limited due to the absence of a protective shelter.

To examine the behavioral response to distal threats, mice were exposed to a sweeping visual stimulus (De Franceschi et al., 2016). Consistent with previous results, mice exposed to sweeping stimuli engaged in passive avoidance strategies, namely immobility to avoid detection (Figure 2A). At the population level, mice showed a robust increase in immobility in the first 20 seconds after the onset of the sweeping stimulus (Figure 2B; baseline: 12.7 ± 3.4% immobility; response: 34.2 ± 3.3% immobility; paired t-test: p < 0.0003, t = 4.448, df = 19, n = 20 mice). We observed no differences in stimulus-evoked immobility across sex (Figure 2C; males: 35.5 ± 6.0% immobility; females: 33.0 ± 2.9% immobility; unpaired t-test: p = 0.71, t = 0.376, df = 18). Despite the lack of sex differences in overall freezing responses, males showed a significantly more variable response to innate threat, contrary to behavioral observations in response to conditioned threat (Gruene et al., 2015; F test: p = 0.04, F = 4.33, df = 9).

**Figure 2:**
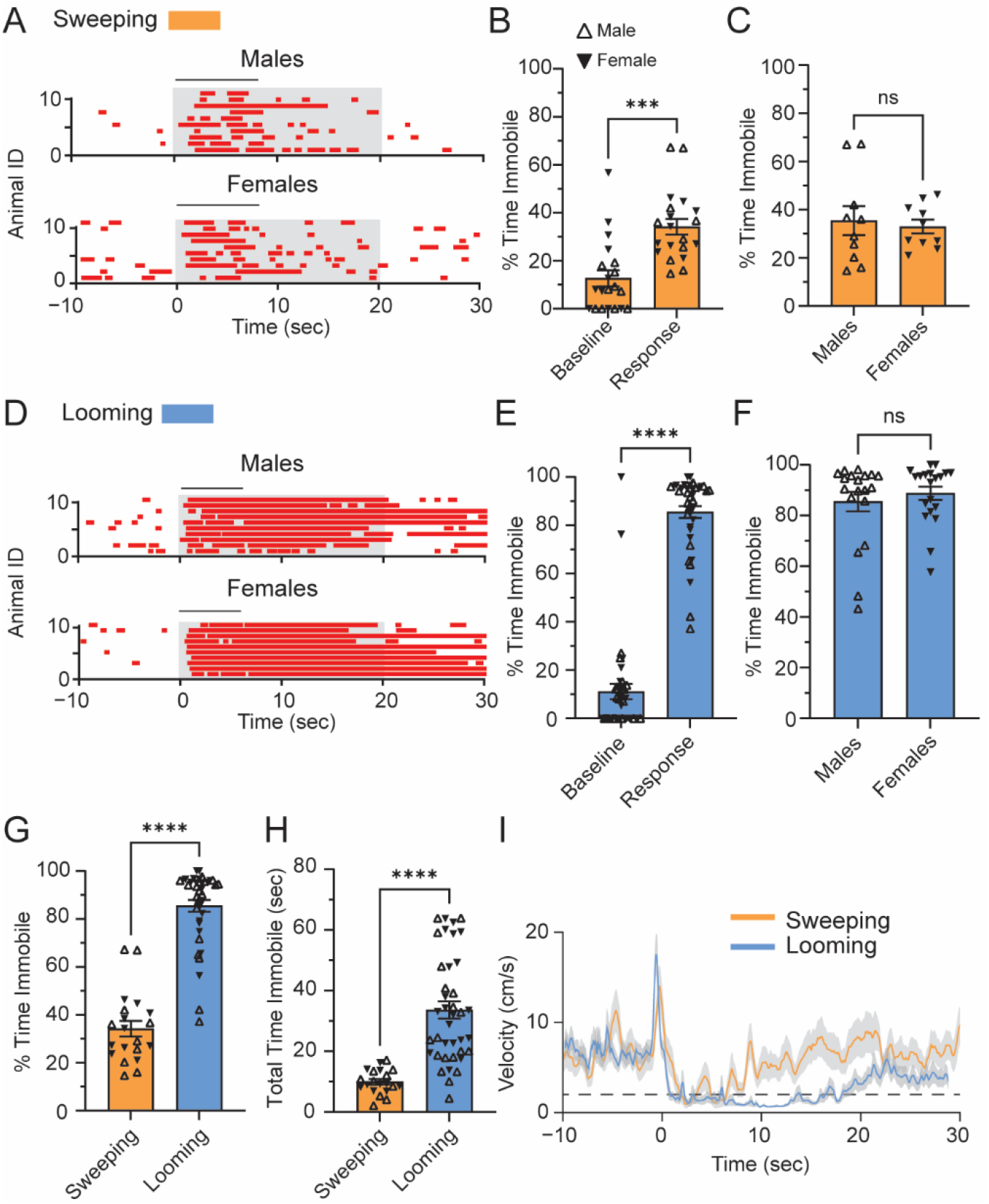
Freezing responses are elicited by looming and sweeping visual stimuli. (A) Stimulus evoked immobility responses triggered by sweeping visual stimuli in male and female mice. Periods of immobility (red, velocity < 2 cm/sec) are indicated for each animal. For clarity, the analysis window (grey box) and stimulus duration (black bar) are also indicated. (B) Sweeping visual stimuli elicit a significant increase in immobility in the 20 seconds after stimulus presentation. (C) There was no difference in the immobility responses across sex. (D) Immobility responses to looming visual stimuli. (E,F) Looming stimuli significantly increased the percent time immobile (E) with no difference observed across sex (F). (G, H) Looming stimuli elicited more robust freezing behaviors, as measured by the percent time freezing (G) and the total time immobile (H). (I) Average velocity as a function of time for all animals for sweeping (left) and looming (right) stimuli. Looming stimulus: n=19 mice, 9 males, 10 females; Sweeping stimulus: n=20 mice, 10 males, 10 females.

In the absence of a shelter, we reasoned that looming threats may result in one of two behavioral responses. One possibility is that mice will engage in an un-directed darting strategy, as has been observed, preferentially in females, in response to conditioned footshocks (Gruene et al., 2015).

Alternatively, mice may respond to looming threats with increased immobility (De Franceschi et al., 2016), in an attempt to avoid detection. In a separate cohort of mice, animals responded to looming threats with a robust increase in immobility (Figure 2D,E; baseline: 16.8 ± 3.4% immobility; response: 87.7 ± 2.3% immobility; paired t-test: p < 0.0001, t = 15.75, df = 38), which was not significantly different between males and females (Figure 2F; males: 85.5 ± 3.8% immobility; females: 88.8 ± 2.6% immobility; unpaired t-test: p = 0.48, t = 0.712, df = 37). These data suggest that under behavioral conditions in which flight to a shelter is not possible, male and female mice similarly engage in defensive *immobility* to both proximal and distal threats.

Interestingly, when directly comparing behavior across stimuli, we observed significantly more immobility to the looming as compared to the sweeping stimulus (Figure 2G; sweeping: 34.2 ± 3.3% immobility, n = 20 mice; looming: 87.2 ± 2.3% immobility; unpaired t-test: p < 0.0001, t = 13.48, df = 57). Considering that the % immobility measurement only considers the 20 second window after stimulus presentation, we additionally calculated the total time each animal engaged in immobility. This measurement accounts for immobility across the entire behavioral trial (i.e. not limited to the first 20 seconds). This measurement similarly showed comparatively more immobility in response to the looming stimulus, suggesting that the differences in immobility were not restricted to the 20 seconds immediately following the stimulus (Figure 2H; sweeping: 9.9 ± 0.9 seconds, n = 20 mice; looming: 35.0 ± 2.7 seconds, n = 19 mice, unpaired t-test: p < 0.0001, t = 6.46, df = 57).

To more fully capture the dynamic responses of animals to threatening stimuli, we averaged the velocity across animals thereby avoiding the categorical classification of animal behavior. In agreement with the above data, looming stimuli resulted in a comparatively more robust and prolonged decrease in velocity (Figure 2I). Interestingly, however, in response to both looming and sweeping stimuli, mice showed a transient elevation in velocity prior to freezing (looming stimulus: 31.3 ± 4.6 cm/s; sweeping stimulus: 20.1 ± 2.8 cm/s, n = 20 mice) which was not significantly different across stimulus type (unpaired t-test: p = 0.10, t = 1.657, df = 57). This transient increase in velocity may represent an initial orienting behavior to assess if an active defensive strategy, such as flight, is a viable response (Evans et al., 2018). These data indicate that in response to both proximal and distal innate threats, mice engage in the optimal behavioral strategy available (in this case immobility), and that they encode threat imminence (i.e. threat proximity) with a significant increase in behavioral vigor.

### Responses to repeated visual stimuli

Fear responses, including innate defensive behaviors, may be modulated by a variety of intrinsic and extrinsic factors such as environmental context, threat history, and internal state (De Franceschi et al., 2016; Hassien et al., 2020; Tafreshiha et al., 2021; Yilmaz & Meister, 2013). We therefore sought to test whether repeated presentation of threatening stimuli (i.e. differential threat history) may alter fear responses at short and long-time scales. We considered two possibilities: either mice will show stable immobility across trials or will show a progressive habituation (Rankin et al., 2009), which may be differentially engaged depending on the proximity (i.e. imminence) of the threat.

To begin to test this, in a single behavioral session, we exposed mice to a series of three repeated sweeping or looming stimuli, each separated by ∼5 minutes. Contrary to our hypothesis, repeated presentation of both stimuli resulted in a gradual decrease in immobility across trials (Figure 3 A, B). In mice exposed to sweeping stimuli, a repeated measures one-way ANOVA revealed a significant decrease in freezing responses across trials (Figure 3C; RM one-way ANOVA: p = 0.0007, F(1.765, 33.54) = 9.869, n = 20 mice), indicative of fear suppression or habituation of immobility. Similarly, looming stimuli elicited robust habituation across trials (Figure 3D; RM one-way ANOVA: p = < 0.0001, F(1.634, 29.40) = 15.79). To measure the total degree of habituation, we used a post-hoc multiple comparison test to compare the freezing response on trial 1 and trial 3 in mice exposed to either looming or sweeping threats. Both stimuli resulted in significant habituation on trial 3 (sweeping: trial 1 vs trial 3: Tukey multiple comparison t-test: p = 0.0015; looming: trial 1 vs trial 3: Tukey multiple comparison t-test: p < 0.0001). To facilitate more direct comparisons *across* both datasets, we calculated the habituation index as a normalized metric of total habituation in each animal, thereby correcting for differences in overall freezing observed across stimuli. These data indicate that the degree of habituation did not differ as a function of stimulus type (Figure 3E; sweeping: habituation index: 0.57±0.08; looming: habituation index: 0.59±0.08; unpaired t-test: p = 0.83 t = 0.214, df = 37). Additionally, we compared the rate of habituation by normalizing immobility within each animal to their response on stimulus 1. Our results indicate that in naïve mice, the rate of habituation across trials was well-fit by a linear regression. Furthermore, the overall rate of habituation did not significantly differ between looming (Figure 3F; slope = -0.20 ± 0.05) and sweeping (slope = -0.22 ± 0.03; p = 0.91, F(1,113) = 0.012). Taken together, these results indicate that freezing behaviors habituate across repeated trials, regardless of the stimulus. Furthermore, the observed linear decrease in immobility occurs on a relatively rapid timescale, suggestive of rapid circuit-level changes in sensorimotor processing.

**Figure 3:**
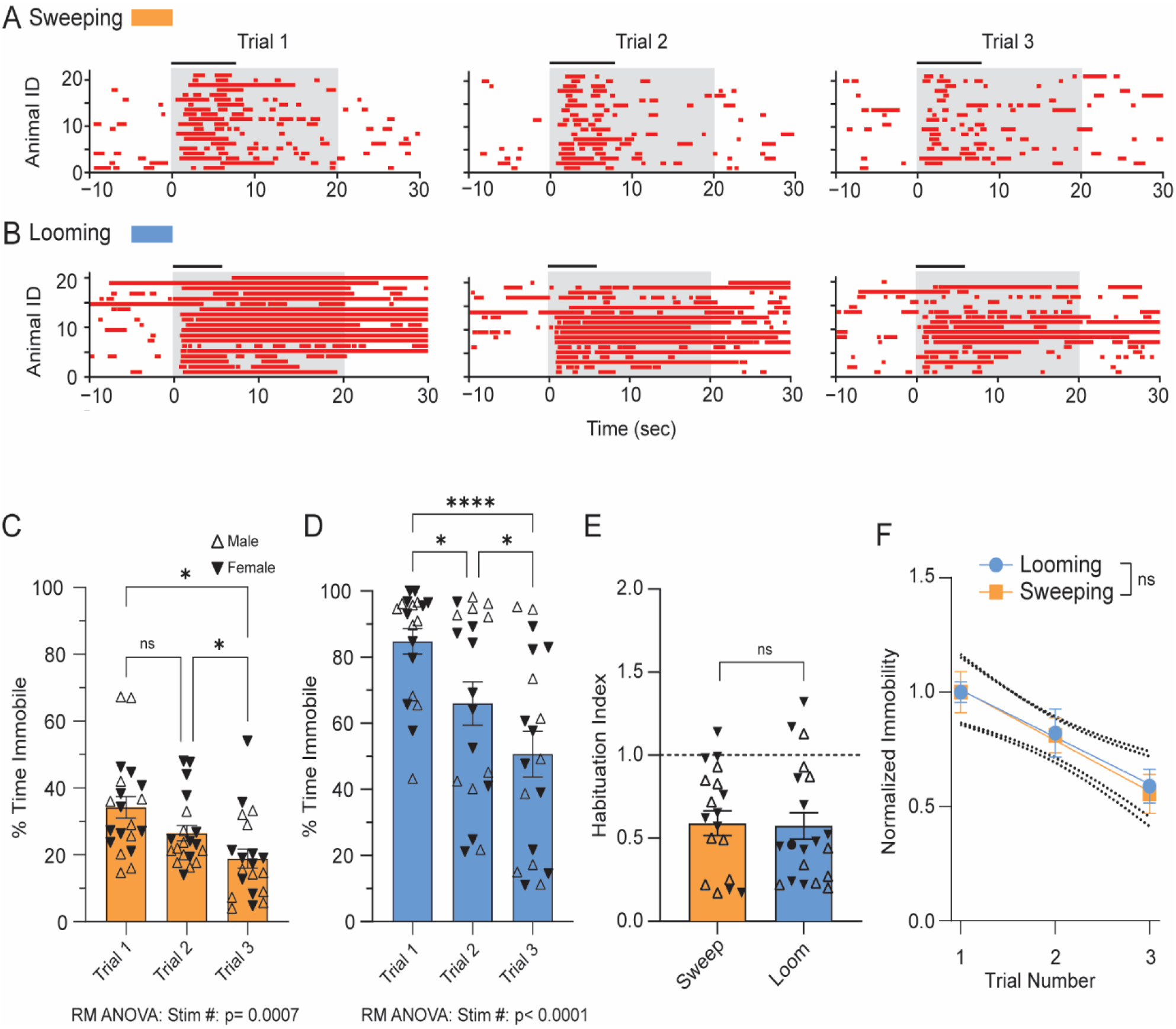
Repeated threat presentation results in robust habituation of freezing behavior. (A, B) Repeated presentation of sweeping (A) or looming (B) visual stimuli at inter-trial intervals as short as 5 minutes results in a gradual reduction in immobility across trials. (C, D) Quantification of immobility across trials for sweeping (C) and looming (D) stimuli. (E, F) Repeated presentation of sweeping and looming stimuli results in equivalent levels of total habituation (E) as well as similar rates of habituation across trials. (F) Normalized immobility responses across trials reveals linear change in immobility. Looming stimulus: n=19 mice, 9 males, 10 females; Sweeping stimulus: n=20 mice, 10 males, 10 females.

We next wondered whether the observed habituation was dependent on the relatively short time between threatening stimuli. To test this, mice were exposed to an identical stimulus paradigm (i.e. three presentations of a looming visual stimulus) but each trial was separated by 24 hours rather than 5 minutes. If habituation resulted from a reduction in threat salience at relatively short time scales, then we would predict more stable immobility across repeated trials at longer time scales (i.e. 24 hours). Contrary to this prediction, looming stimuli presented at intervals of 24 hours, resulted in an enhanced habituation across trials (Figure 4A; RM one-way ANOVA: F(1.631, 30.99) = 38.06, p < 0.0001, n = 20 mice). Interestingly, increasing the interval between stimuli appeared to alter the overall pattern of habituation across trials. To quantify these changes, we first used a two-way repeated measures ANOVA to compare the overall pattern of habituation in mice exposed to looming stimuli separated by 5 minutes and 24 hours. As expected, there was a significant main effect of stimulus number (p < 0.0001, F(1.913, 70.76) = 4.05). Additionally, there was a significant interaction between stimulus number and inter-trial interval (p = 0.02, F(2,74) = 4.05), suggesting that the overall pattern of habituation differed as a function of inter-trial interval. The observed change in habituation pattern resulted from a non-linear change in habituation across trials. More specifically, at 24-hour inter-trial intervals, we observed a significant decrease in freezing between trials 1 and 2 (Tukey’s multiple comparisons test: adjusted p: <0.0001) but no difference in immobility between trials 2 and 3 (Tukey’s multiple comparisons test: adjusted p: 0.841). This pattern suggests that habituation is accelerated at longer inter-trial intervals. To quantify the accelerated habituation, we compared the change in freezing between trials 1 and 2 at both inter-trial intervals. Consistent with the observation of accelerated habituation, the change in the freezing duration between trials 1 and 2 was significantly larger at 24 hours (-22.24 ± 2.5%, n = 20 mice) than at 5 minutes (12.25 ± 2.9%, n = 19 mice; unpaired t-test: p = 0.012, t=2.617, df = 37). However, despite the accelerated habituation, the overall degree of habituation measured on trial 3 was similar across inter-trial intervals (Figure 4C; 5 minute interval: habituation index: 0.60 ± 0.08; 24 hour interval: habituation index: 0.47 ± 0.05; unpaired t-test: p = 0.19, t = 1.336, df = 37). Together, these data indicate that the overall rate of habituation depends on the interval between trials, however, the total degree of habituation is consistent regardless of inter-trial interval.

**Figure 4:**
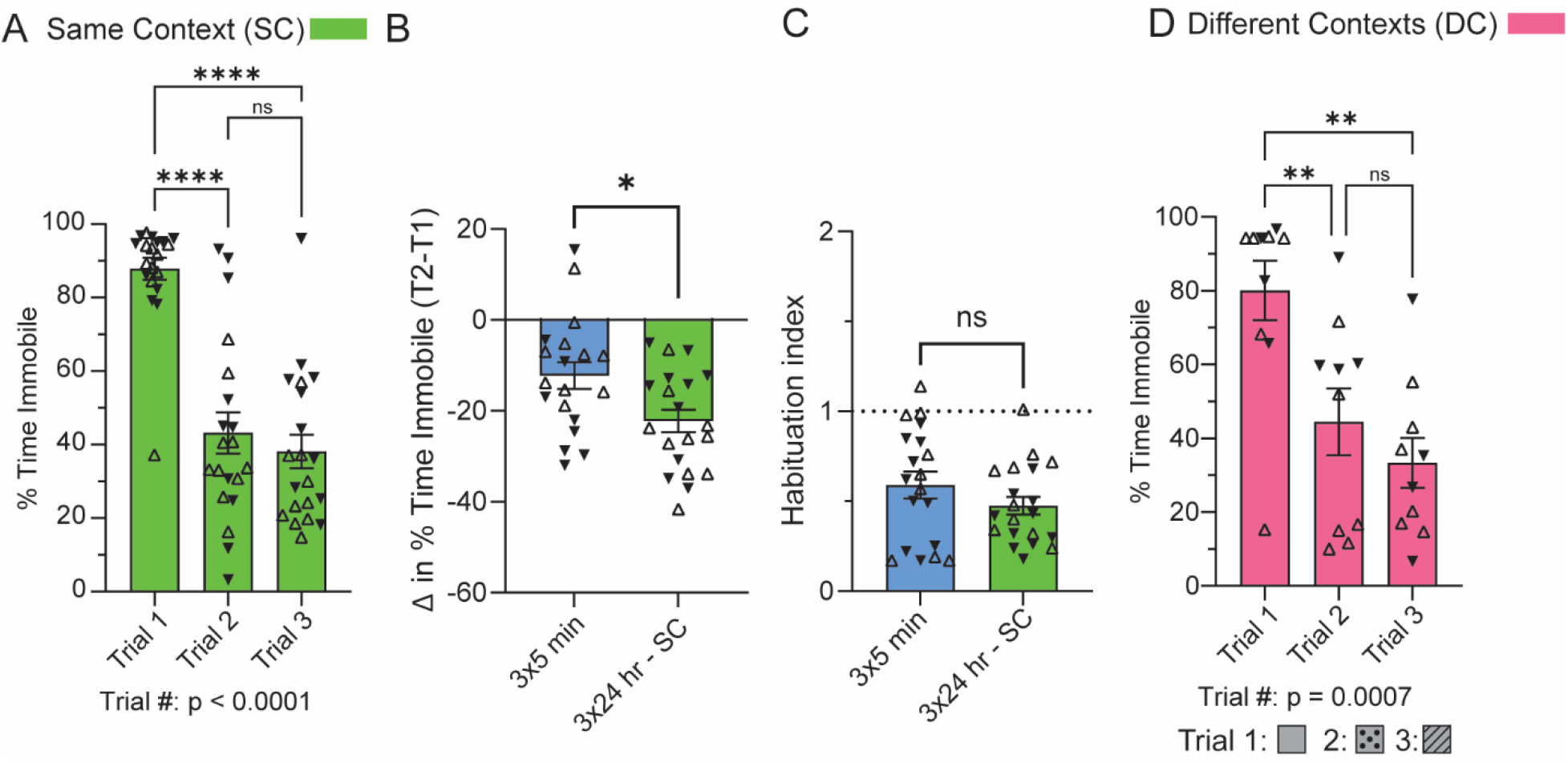
Accelerated pattern of habituation when the looming visual stimuli was separated by 24 hour intertrial intervals. (A) Immobility response to repeated looming visual stimuli presented at 24 hour inter-trial intervals in a behavioral chamber with identical contextual cues across trials. (B) Comparing the change in immobility from the first and second stimulus presentation (Trial 2-Trial 1) between animals that received the repeated looming visual stimuli with ∼5 minute inter-trial intervals (Blue) or ∼24 hour inter-trial intervals (Green) revealed an accelerated rate of habituation in 3×24 animals. (C) Normalized rate of habituation in 3×5 and 3×24 animals showed similar rates of overall innate fear behavioral habituation. (D) Immobility responses at inter-trial intervals of 24 hours in distinct behavioral contexts was similar to those observed in the same context. 3×5 min: n=19 mice, 9 males, 10 females; 3×24 SC: n=20 mice, 10 males, 10 females; 3×24 DC: n=10 mice, 5 males, 5 females.

The observed accelerated habituation at longer inter-trial intervals could be explained, in part, by contextual fear learning extinction (Maren et al., 2013), as stimuli were presented in identical contexts across all three days. To explicitly test whether the accelerated habituation was mediated by contextual cues, we repeated the experiment in a new cohort of mice, in which the context was varied between trials, with an inter-trial interval of 24 hours. Under these experimental conditions, the overall pattern of habituation was similar to that observed at 24 hour inter-trial intervals in a single context. More specifically, we observed non-linear, accelerated habituation, similar to that observed in a single context (Figure 4D; RM one-way ANOVA: p = 0.0007, F(1.912, 17.21) = 11.7), which was not significantly different than that observed in a single context (two-way ANOVA interaction: p = 0.61 F(2,56) = 0.505). Together, these results indicate that contextual cues were insufficient to explain the accelerated habituation at extended inter-trial intervals. Furthermore, our results suggest that the neural mechanisms underlying behavioral habituation may be distinct at short and long-time scales.

### Freezing responses are diminished in novel contexts

We next sought to determine whether innate freezing differs in mice that have been exposed to the behavioral arena through familiarization versus mice in a completely new environment. We reasoned that in novel environments, mice may show increased behavioral vigilance resulting in increased fear responses, as they explore the novel environment. Alternatively, mice may disregard potentially threatening stimuli as they familiarize themselves with their surroundings, which would result in a decreased freezing behavior. To test this, mice were not familiarized with the open field chamber prior to behavioral testing. To test whether environmental novelty impacted behavioral habituation, we presented mice with 3 looming stimuli separated by 5 minutes, as in previous datasets. In response to the initial stimulus presentation, mice in novel environment showed a small yet statistically significant immobility response (Figure 5A; baseline: 15.5 ± 3.6% time immobile; response: 28.2 ± 4% time immobile, paired Student’s t-test: p = 0.01, t = 3.1, df = 19). However, when compared to a separate cohort of mice that underwent chamber familiarization (as in previous cohorts), mice exposed to looming stimuli in novel environments showed a significantly attenuated fear response (Figure 5B; novel: 28.2 ± 4 % time immobile; familiar: 72.2 ± 5.2% time immobile; unpaired Student’s t-test: p < 0.0001, t = 6.8, df = 38). Similar results were obtained when comparing the total time immobile in the minute following stimulus presentation (Figure 5C; novel: 9.6 ± 1.6 seconds; familiar: 23.7 ± 2.2 seconds; unpaired Student’s t-test: p < 0.0001, t = 5.13, df = 38).

**Figure 5:**
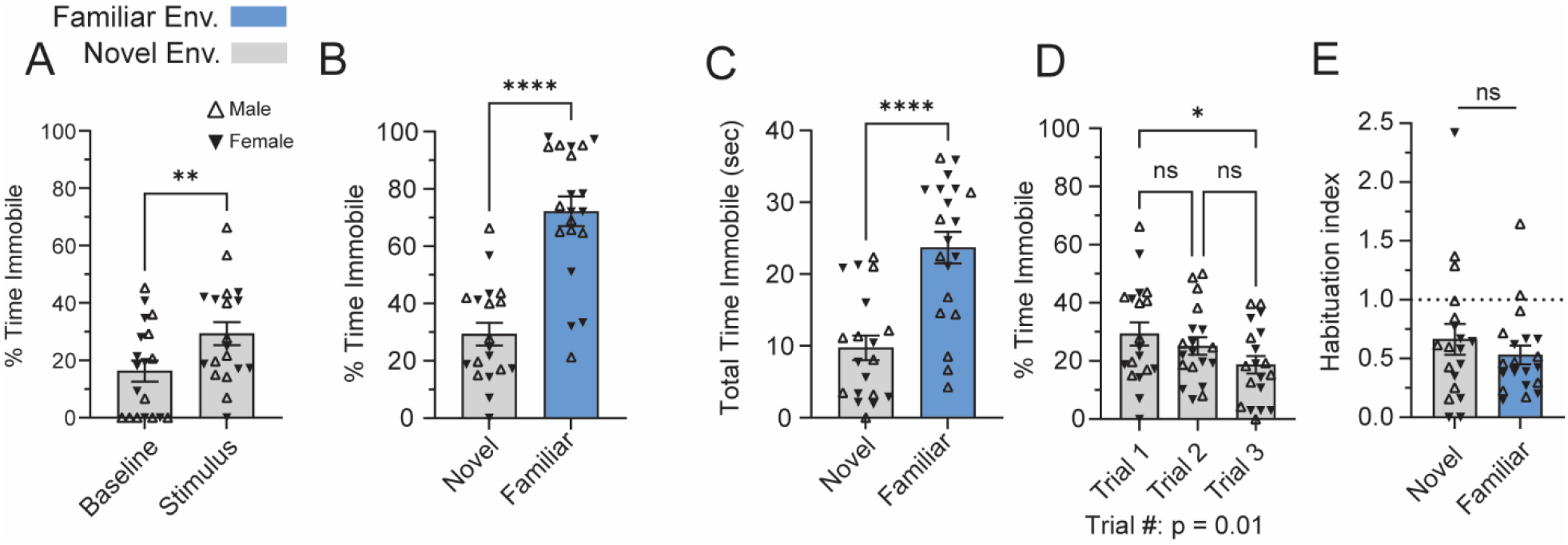
Freezing response in a Familiar (blue) and novel (light grey) environment. (A) Immobility response to a looming visual stimulus in mice that were not familiarized with the testing environment before behavioral testing. (B) Comparison of percent time immobile between familiarized (Blue) and unfamiliarized (Grey) animals in response to a looming visual threat. (C) Total time immobile in seconds during the duration of video recording after looming stimulus onset, exceeding the 20 second analysis window used to calculate percent time immobile. (D) Repeated presentations of the looming visual stimulus at inter-trial intervals of 5 minutes results in a reduced freezing response in mice within a novel environment. (E) Comparison of normalized habituation in mice exposed to looming stimuli under novel and familiar environmental conditions. Familiarized: n=20 mice, 10 males, 10 females; Novel: n=19 mice, 9 males, 10 females.

Despite the lower overall freezing behavior in response to a single looming stimulus, mice in novel environments still showed significant habituation across repeated trials, as there was a significant main effect of trial number on freezing responses in a repeated measure ANOVA (Figure 5D; RM one-way ANOVA: F(1.6, 31.05) = 3.7, p = 0.07; trial 1 vs. trial 3: Tukey post-hoc comparison: p = 0.015). Compared to mice in familiar environments, the rate of habituation was delayed in the novel environment, as there was no significant difference in freezing across trials 1 and 2 (Figure 5C; Tukey’s post-hoc comparison: p = 0.47). However, the total degree of habituation was not significantly different in mice exposed to looming stimuli novel versus familiar environments (Figure 5E; novel: 0.66 ± 0.57; familiar: 0.53 ± 0.36; unpaired Student’s t-test: p = 0.38, t = 0.9, df = 37, n = 19 mice (novel), 20 mice (familiar), one mouse was removed from analysis in the novel condition as an outlier exceeding >3 standard deviations from the population mean). These results indicate that mice in novel environments demonstrate both a reduced overall fear response and a delayed habituation profile across repeated trials.

### Effects of acute stress on innate freezing

In addition to extrinsic factors, such as the nature of the visual stimulus, environmental context, or context familiarity, fear responses can also be modulated by physiological factors such as internal state. For example, in conditioned fear paradigms, exposure to intensely threatening acute stress has been shown to sensitize both associative and non-associative fear learning (Hassien et al., 2020; Perusini et al., 2016; Rau et al., 2005; Rau & Fanselow, 2009). We therefore sought to test whether similar acute stress paradigms resulted in changes in *innate* fear responses. To do this, mice were exposed to a set of four unconditioned footshocks (amplitude: 1 mA, duration: 2 seconds, inter-trial interval: randomly applied between 60-90 seconds, as in (Hassien et al., 2020)) in a distinct behavioral context either 1 hour (Figure 6A) or 24 hours (Figure 6B) prior to exposure to looming threats. We first compared the immobility response on the first of three visual stimuli and found no significant effect of acute stress on initial innate fear responses (Figure 6C; Ordinary one-way ANOVA: F(2,54) = 2.06, p = 0.14).

**Figure 6:**
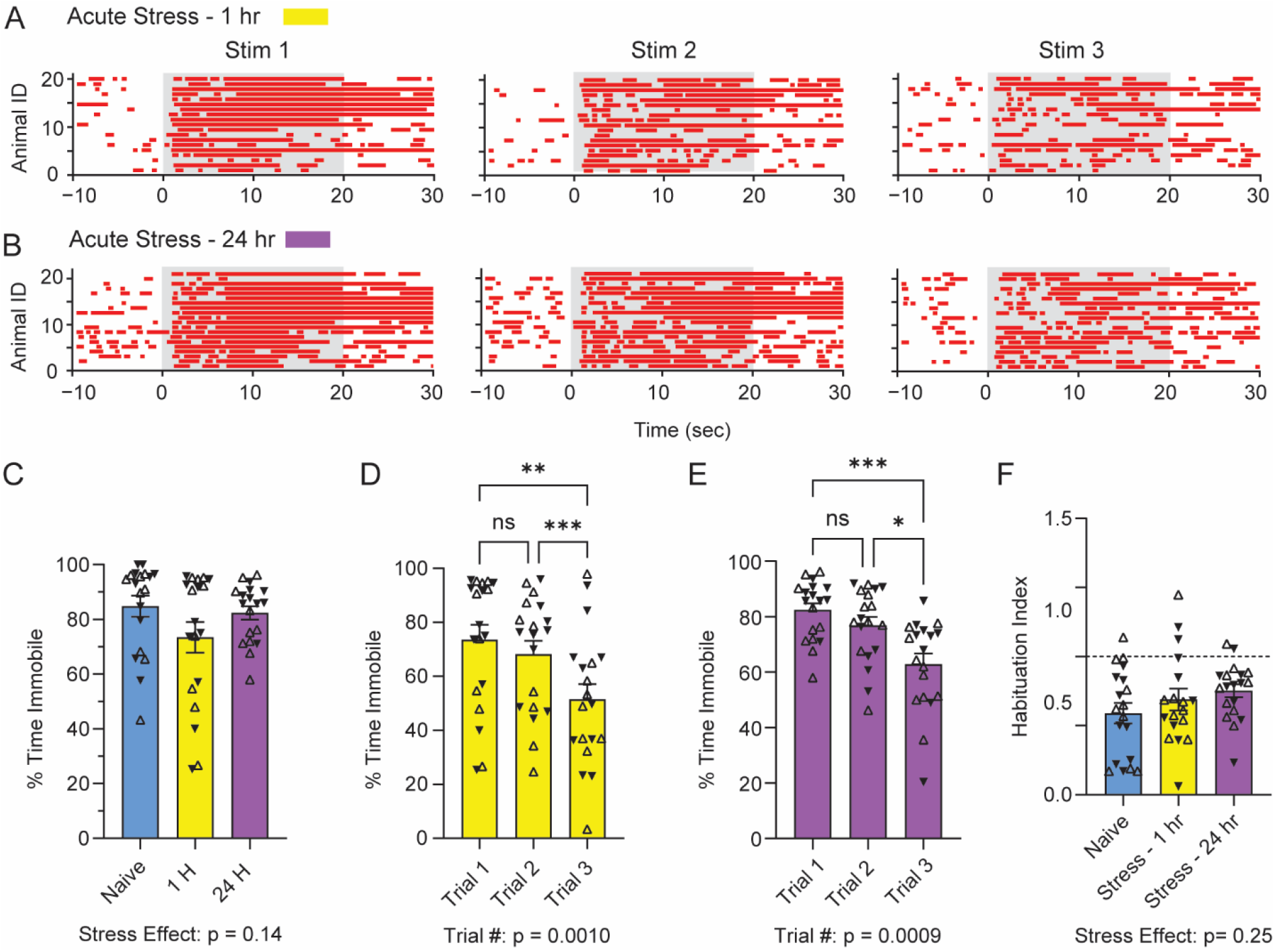
Acute stress effects on innate fear behaviors. (A,B) Immobility responses (red, velocity <2 cm/sec) triggered by Looming visual stimuli in male and female mice. Mice were exposed to acute footshock stress either 1 hour (A) or 24 hours (B) before exposure to innate fear paradigm. (C) Comparison of the freezing response on trial 1 across naïve and stressed animals. (D, E) Quantification of freezing behavior across trials in mice exposed to footshock stress 1 hour (D) or 24 hours (E) before the innate fear paradigm. (F) Comparison of the overall level of habituation across all three cohorts (naïve and stressed). Naïve: n=19 mice, 9 males, 10 females; Acute stress 1 hour: n=19 mice, 9 males, 10 females; Acute stress 24 hours: n=19 mice, 10 males, 9 females (one mouse was removed from analysis in the novel condition as an outlier exceeding >2.5 standard deviations from the population mean).

We next examined the pattern of habituation across trials, which together demonstrated that the rate of innate fear habituation was significantly delayed at 1 hour and 24 hours after acute stress. At both time points, a repeated measure one-way ANOVA revealed a significant main effect of trial number on freezing (Figure 6D-E 1 hour post-stress: RM one-way ANOVA: F(1.505, 27.09) = 10.62, p = 0.001; 24 hours post-stress: F(1.575, 28.36) = 10.46, p = 0.0009).

Furthermore, in both datasets mice showed a significant decrease in freezing when comparing trials 1 and 3 (Figure 6D-E; 1 hour post-stress: Tukey post-hoc comparison: adjusted p = 0.0062; Figure 6E; 24 hours post-stress: Tukey post-hoc comparison: adjusted p = 0.0005), indicating intact habituation across trials. However, there was no statistically significant difference in freezing between trials 1 and 2 (1 hour post-stress: Tukey post-hoc comparison: adjusted p = 0.051; 24 hours post-stress: Tukey post-hoc comparison: adjusted p = 0.26), suggesting that the habituation was significantly delayed. Finally, we compared the overall degree of habituation across naïve (unstressed) animals and animals exposed to stress and found no significant difference across all three datasets (Figure 6F; ordinary one-way ANOVA: F(2,53) = 1.514, p = 0.23) indicating that stress does not significantly impact the overall degree of innate fear habituation. Together, these data suggest that changes in internal state, such as those following an unpredicted stressor, can significantly delay the rate of innate fear habituation, but have little effect on overall freezing levels or the total degree of habituation across trials.

## Discussion

Here, we set out to determine whether and how threat imminence is encoded in behavioral action following innate predator threats, specifically under conditions in which the available defensive reactions are limited. Our results indicate that under experimental conditions in which escape behaviors are disfavored, namely in the absence of a protective shelter, mice responded to both looming and sweeping stimuli with robust immobility. Both male and female mice showed comparatively increased immobility to the looming stimulus, which, ethologically, represents a more proximal threat. These results suggest that mice encode threat imminence not only in behavioral action selection, but also in response vigor. Furthermore, our results indicate that in response to repeated presentation of looming and sweeping threats, mice consistently reduce immobility across trials, reflecting habituation. However, the overall pattern of fear suppression differed across experimental manipulations, suggesting that innate fear suppression is subject to modulation by environmental (extrinsic) and physiological (intrinsic) factors.

### Threat imminence theory and species-specific defense reactions

One predominant and unifying theory of innate (and conditioned) fear, is that fear serves to restrict the behavioral repertoire of prey animals to a circumscribed, species-specific set of defensive reactions designed to promote survival (Bolles, 1970; Crawford & Masterson, 1982; Fanselow, 2018; Fanselow & Lester, 1988). This theory has been further expanded upon with the development of the threat imminence theory (R. J. Blanchard & Blanchard, 1971; Bolles & Fanselow, 1980; Fanselow & Lester, 1988), which postulates that as threats shift from distal to proximal, animals respond with distinct, ethologically appropriate behavioral strategies, likely mediated by distinct neural circuits (Deng et al., 2016; Fanselow, 1991, 1994; Gale & Murphy, 2014; Shang et al., 2018). For example, a distal aerial predator simply flying overhead may initiate a ‘passive coping strategy’ such as freezing to avoid detection whereas an approaching aerial predator will elicit more active avoidance strategies, such as flight, to evade capture (De Franceschi et al., 2016; Tafreshiha et al., 2021; Yilmaz & Meister, 2013). It is worth noting that the dichotomy of passive vs. active avoidance strategies is largely one of semantic convenience, as ‘passive coping strategies’ are intentionally engaged and often involve complex, whole-body motor coordination, and are therefore not truly passive. Despite this qualification, the threat imminence theory is an influential model to describe how animals appropriately regulate or adapt their defensive strategy in response to varied environmental threats.

Innate fear responses have evolved from an evolutionary pressure to avoid predation risk, resulting in species-specific defensive reactions that are tuned to specific ecological niches. For example, closely related species of *Peromyscus* mice engage with identical predator threats using distinct behavioral strategies that are ethologically matched to their evolutionary history (Baier et al., 2023; Hirsch & Bolles, 1980). In addition to the diversity of behavioral strategies observed, prey species must also show a high degree of behavioral flexibility to select the optimal defensive strategy depending on external conditions such as proximity to a nest or other environmental factors (Campagner et al., 2022; Evans et al., 2018; Lefler et al., 2020; Vale et al., 2017). Although the threat imminence theory provides a powerful general framework to describe how threat proximity influences defensive strategies, understanding how threat imminence is encoded in other behavioral metrics of fear, such as behavioral vigor, has been much more difficult to assess. This difficulty arises, in part, due to the inherent variability in behavioral strategy employed by mice exposed to proximal or distal threats.

To facilitate a more direct comparison of behavioral vigor across proximal and distal threats, we designed an innate fear behavioral paradigm in which both looming and sweeping visual stimuli were presented to mice in a chamber lacking a protective shelter. Our results indicate, consistent with previous literature (De Franceschi et al., 2016; Yilmaz & Meister, 2013), that under such conditions mice engage in freezing behaviors in response to both looming and sweeping visual stimuli. Here we have directly compared responses to looming and sweeping visual threats, which is consistent with previous literature in experiments without a shelter (De Franceschi et al., 2016). Our results expand on the previous findings by suggesting that mice still encode threat proximity in the behavioral vigor with which they respond, as more proximal threats elicited more robust (longer bouts of) defensive freezing.

It is worth noting that freezing in response to a looming predator may be considered ethologically counter-intuitive as immobility in the face of an approaching predator may increase the risk of predation. However, our data indicates that the strict behavioral hierarchy proposed by the threat imminence theory is incomplete. In the absence of a protective shelter, mice engage in defensive immobility, presumably to reduce the odds of detection while predators make fine-scale adjustments to their attack trajectory (i.e. not all attacks are ballistic). In fact, our data suggest that proximal threats result in increased behavioral vigor, as demonstrated by the increased duration of immobility. In such cases (i.e. the absence of a shelter) the ‘optimal’ strategy may be prolonged freezing to increase the probability that the predation attempt is unsuccessful. Conversely, more distal threats, by definition, involve less risk, which results in reduced behavioral vigor. Overall, our results reinforce the concept that fear limits the available behavioral repertoire and further reinforce the threat imminence theory, by suggesting that in addition to threat proximity influencing behavioral choice, it is also encoded in response vigor.

### Behavioral flexibility is critical for appropriate fear responses

In addition to ethological pressures to select the optimal behavioral strategy to avoid predation, animals must also be able to assess and adjust their defensive responses. As such, properly regulated fear responses require animals to both respond appropriately to threats in the environment while simultaneously adjusting fear responses to non-threatening stimuli (Tafreshiha et al., 2021). Consistent with this view, our data demonstrate that repeated presentation of looming and/or sweeping visual stimuli resulted in rapid decreases in freezing behavior across repeated trials, which could be engaged with inter-trial intervals as short at 5 minutes. Such habituation may be ethologically adaptive, as it allows mice to re-evaluate whether specific sensory inputs reflect acute threats in the environment. The relatively short timescales at which habituation was observed suggests that the neural mechanisms underlying such habituation may be mediated by rapid changes in synaptic integration in central fear circuits. For example, emerging evidence suggests that circuits in the periaqueductal gray, a central hub for generating fear behaviors (D. C. Blanchard & Blanchard, 2008; Koutsikou et al., 2015; C. Silva & McNaughton, 2019), may be modulated by numerous upstream circuits including the cerebellum (Vaaga et al., 2020), in a direction predicted to reduce freezing responses.

The behavioral habituation observed in response to repeated innate visual threats is reminiscent of extinction learning in instrumental or Pavlovian conditioning paradigms (Maren et al., 2013), although the underlying neural mechanisms may be entirely distinct. In conditioned paradigms, extinction learning occurs when the reinforcing unconditioned stimulus (i.e. foot shock) is no longer presented in conjunction with the conditioned stimulus (i.e. tone) (Bouton et al., 2021; Delamater & Westbrook, 2014; Quirk & Mueller, 2008). There has been great interest in understanding the behavioral and neural mechanisms underlying extinction, as it allows animals to adjust their behavior to novel environments. Our results demonstrate that numerous intrinsic and extrinsic factors (such as inter-trial interval, chamber familiarity and exposure to acute stressors) alter the rate of habituation in an innate fear paradigm across repeated trials.

### Impact of internal state on innate fear responses

Of particular interest is the observed impact of acute stress on innate fear responsivity both at 1 hour and 24 hours after stress exposure. Previous work has demonstrated that acute, unpredicted stress significantly increases both associative and non-associative fear responses in both mice and rats (Hassien et al., 2020; Perusini et al., 2016; Perusini & Fanselow, 2015; Rau et al., 2005; Rau & Fanselow, 2009), which has been proposed as a fundamental model of post-traumatic stress disorder (PTSD). Our work suggests that while acute stress does not significantly impact the fear response on the first trial, it does significantly delay fear habituation. Functionally, the delayed habituation represents an enhanced fear state – in which mice maintain robust freezing responses for a prolonged period. This enhanced fear state is consistent with the observed effects on fear learning, which are enhanced following similar acute stress protocols. The increase in fear state across 24 hours suggests that the effects are not mediated directly by enhanced circulating corticosterone, but rather may involve long-term synaptic remodeling in innate fear circuitry, including the periaqueductal gray (Myers et al., 2014). Interestingly, NMDA receptor activation is required for acquisition of associative fear memory in the stress context (Rau et al., 2005), whereas circulating corticosterone is required for both increased associative fear memory and stress-enhanced fear learning (Perusini & Fanselow, 2015). Finally, our data support the accumulating evidence in favor of the stress-enhanced fear learning paradigm as a model for PTSD (Hassien et al., 2020; Perusini & Fanselow, 2015; Rau et al., 2005), as the delayed habituation profile observed at 24 hours post-stress exposure likely represents a unique form of delayed fear extinction, a hallmark clinical feature of PTSD. Future work is therefore needed to resolve the neural mechanisms underlying the stress-induced changes in innate fear responsivity and fear habituation.

## Acknowledgments

We thank Dr. Indira Raman for her support of the initial behavioral work at Northwestern University and for providing helpful comments on the manuscript. We also would like to thank members of the Raman, Myers, and Vaaga labs for helpful discussion and feedback on the project.

